# Computational design of a single-domain antibody that specifically recognizes WT1 peptide-loaded class I MHC

**DOI:** 10.1101/2025.07.24.666704

**Authors:** Tae-Sung Oh, Sang Pil Ahn, Seonghyuk Suh, JooYeon Lee, Yeong-Nam Bae, Hyejin Lee, Yong-Sung Kim, Bo-Seong Jeong, Byung-Ha Oh

**Affiliations:** Therazyne, Inc., Daejeon, Republic of Korea; Department of Biological Sciences, Advanced Institute of Science and Technology, Daejeon, Republic of Korea; Department of Molecular Science and Technology, Advanced College of Bioconvergence Engineering, Ajou University, Suwon, Republic of Korea

**Author notes:** Corresponding authors: (Byung-Ha Oh); (Bo-Seong Jeong).

**Keywords:** Computational antibody design, peptide-loaded class I MHC, TCR-like single-domain antibody, Wilm’s tumor 1 protein

## Abstract

T cell receptor (TCR)-like antibodies that recognize peptide-loaded class I MHC (pMHC) complexes can enable precise targeting of cancer cells, but developing single-domain binders with high specificity and affinity is challenging. Here, we report the computational design and experimental validation of a TCR-like single-domain antibody (sdAb) that specifically recognizes the WT1-derived peptide RMFPNAPYL presented by HLA-A*02:01. Starting from the crystal structure of a Fab antibody bound to RMF/HLA- A02:01, we repurposed the V_H_ domain into a stable, soluble Trastuzumab-derived V_H_ scaffold. The resulting sdAb, RMFsdAb, spans all nine peptide residues and shows no binding to a control pMHC with a different peptide. Its biophysical properties were improved by fusion to human serum albumin domain III (HSA D3), yielding a monodisperse HSA D3-RMFsdAb with 81 nM affinity and specificity for RMF/HLA- A*02:01. We further engineered a bivalent format (RMFsdAb-HSA D3-RMFsdAb), which dramatically increased apparent binding affinity to 0.4 nM. When expressed on a CAR T cell, HSA D3-RMFsdAb functioned as the antigen-recognition domain to selectively activate T cells in the presence of RMF/HLA-A*02:01–positive cells. Our results demonstrate a viable strategy to develop high-specificity, peptide-focused TCR-like sdAbs for pMHC-targeted therapeutics.

## INTRODUCTION

Class I major histocompatibility complex (MHC) molecules are composed of a polymorphic human leukocyte antigen (HLA) heavy chain and a β2-microglobulin (β2M) light chain. Expressed on nearly all nucleated cells, they play a critical role in immune surveillance by enabling the detection of infected or malignant cells. Peptide-loaded MHC (pMHC) antigens on the cell surface are continuously monitored by T-cell receptors (TCRs). When a nonself peptide or an abnormal self-peptide is presented by pMHC, it can activate the responsive T cells, which then target and kill the presenting cell [1]

Developing TCRs specific to target pMHCs is a significant focus in research and pharmaceuticals. Two pMHC-targeting modalities have received clinical approval: Tebentafusp, a TCR-derived bispecific T cell engager (BiTE) targeting HLA-A*02:01- positive melanoma cells presenting a gp100-derived peptide [2] and Afamitresgene autoleucel, an adoptive cell therapy targeting HLA-A*02:01-positive synovial sarcoma cells presenting a MAGE-A4-derived peptide [3].

A TCR-like antibody is a monoclonal antibody that mimics the specificity of a TCR in recognizing pMHCs. These antibodies combine the antigen-binding capacity of an antibody with the ability to recognize a specific peptide epitope presented by MHC molecules. Owing to their stability, versatility, and ease of manufacturability, TCR-like antibodies present a viable alternative to TCR-based therapies. Several TCR-like antibodies targeting pMHCs have been developed, showing promising clinical potential [4–7]. These antibodies cannot be directly used therapeutically unless their target antigens are exceptionally expressed on cancer cells. Instead, they need to be converted into specialized formats like T-cell bispecific antibody [8], bispecific T cell engager (BiTE) [7], chimeric antigen receptor (CAR) [9] or STAR (synthetic TCRs and antigen receptors) in which the TCRα and TCRβ constant regions are fused to the V_L_ and V_H_ domains of a TCR-like antibody, respectively [10]. All of these formats function by activating T cells upon specific engagement with pMHC antigens.

Developing TCR-like antibodies with negligible cross-reactivity is challenging. This is because the size of the real antigen, the peptide displayed on an HLA molecule, is fractional compared to that of MHC. Moreover, the binding surface of a Fab is larger than the surface formed by the bound peptide, often interacting with HLA regions that are similar across subtypes, leading to potential cross-reactivity with non-target pMHC on normal cells. Nanobodies, with their smaller epitope-binding surfaces, could be advantageous TCR-like antibodies in reducing cross-reactivity, and a few TCR-like nanobodies have been developed and tested for effectiveness in the adoptive cell therapy format [11, 12].

Beyond traditional library screening with positive and negative sorting, a strategy for re-engineering preselected TCR-like antibodies was reported to target different pMHC antigens, enabling the isolation of new TCR-like antibodies [13]. In addition, design of non-TCR-like, synthetic scaffolds to target specific pMHCs have also been reported [14, 15].

Wilms’ tumor 1 protein (WT1) is a zinc finger transcription factor highly expressed in various cancers but rarely in normal adult cells [16–19]. WT1-derived peptides are presented by MHC as 9- or 10-amino acids [20, 21]. A 9-mer WT1 peptide (126-RMFPNAPYL-134, ‘RMF’) is presented by the HLA-A*02:01 subtype. TCRs or TCR-like antibodies have been developed that recognize RMF bound to HLA-A*02:01– β2m (for simplicity, denoted as HLA-A*02:01 hereafter) [8, 22–24] and utilized for immunotherapeutic approaches against several cancers such as acute myeloid leukemia [8, 25–27].

To date, no TCRllllike nanobody specific for the RMF/HLAlllA*02:01 complex has been reported. We report the repurposing of the anti-Her2 antibody Trastuzumab’s V_H_ into a TCR-like single-domain antibody (SdAb). This sdAb fully covers the entire length of the bound peptide and exhibits no cross-reactivity to HLAlllA*02:01 loaded with p53- derived peptide.

## RESULTS

### Structural information of an antibody bound to RMF peptide-loaded MHC

A TCR-like antibody against the RMF/HLA-A*02:01 complex, named Q2L, was reported without structural information [24]. We determined the crystal structure of the Q2L-Fab bound to RMF/HLA-A*02:01 at 2.8 Å resolution (Table 1). The asymmetric unit contains two molecules of Q2L-Fab/RMF/HLA-A*02:01. Unexpectedly, the V_L_ domains are separated from their V_H_ counterparts, forming limited intermolecular interactions between the V_L_ domains of the two Fab molecules. These interactions are minor and likely represent crystal contacts rather than stable homodimeric interactions in solution. Furthermore, the V_L_ domain of one Fab molecule is positioned closer to the V_H_ domain of the other Fab than to its own V_H_. Consequently, each RMF/HLA-A02:01 complex interacts with the V_H_ domain of one Fab molecule and the V_L_ domain of the other Fab molecule (Figure 1). The latter interactions, involving His145 and Lys146 of HLA-A02:01 and Ser50 and Leu53 of V_L_, are minor and unlikely to occur in solution.

**Figure 1.**
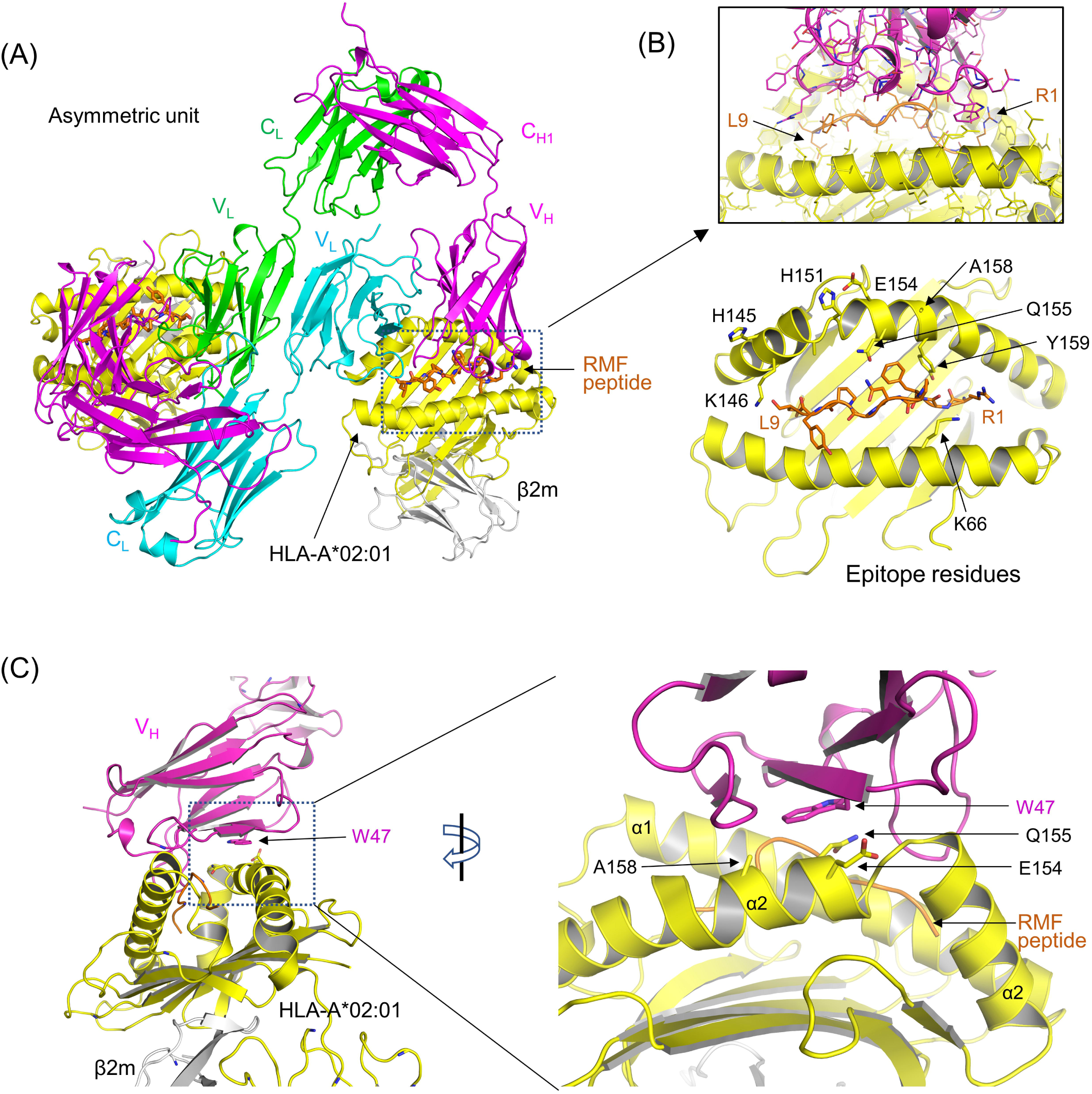
Structural features of Q2L-Fab/RMF/HLA-A*02:01 (A) Overall structure. Two molecules of the complex in the asymmetric unit are displayed. The V_L_ domain of one complex is separated from V_H_ and interacts with the V_L_ domain of the adjacent complex. The enlarged view shows that V_H_ spans the bound RMF peptide. (B) pMHC interaction with Fab. Peptide contacts with V_H_ are shown (*Top*), and pMHC residues within 4 Å of the Fab (six epitope residues) are shown as sticks and labeled (bottom). Also shown are His145 and Lys146 (in grey) from one complex, which interact with the V_L_ domain of the other Fab molecule. (C) Trp47 interaction with HLA-A*02:01. The indole ring of Trp47 in V_H_ sits on helix α2 (labeled) of HLA-A*02:01 and interacts with hydrocarbon atoms of Glu154, Gln155 and Ala158.

**Table 1.**
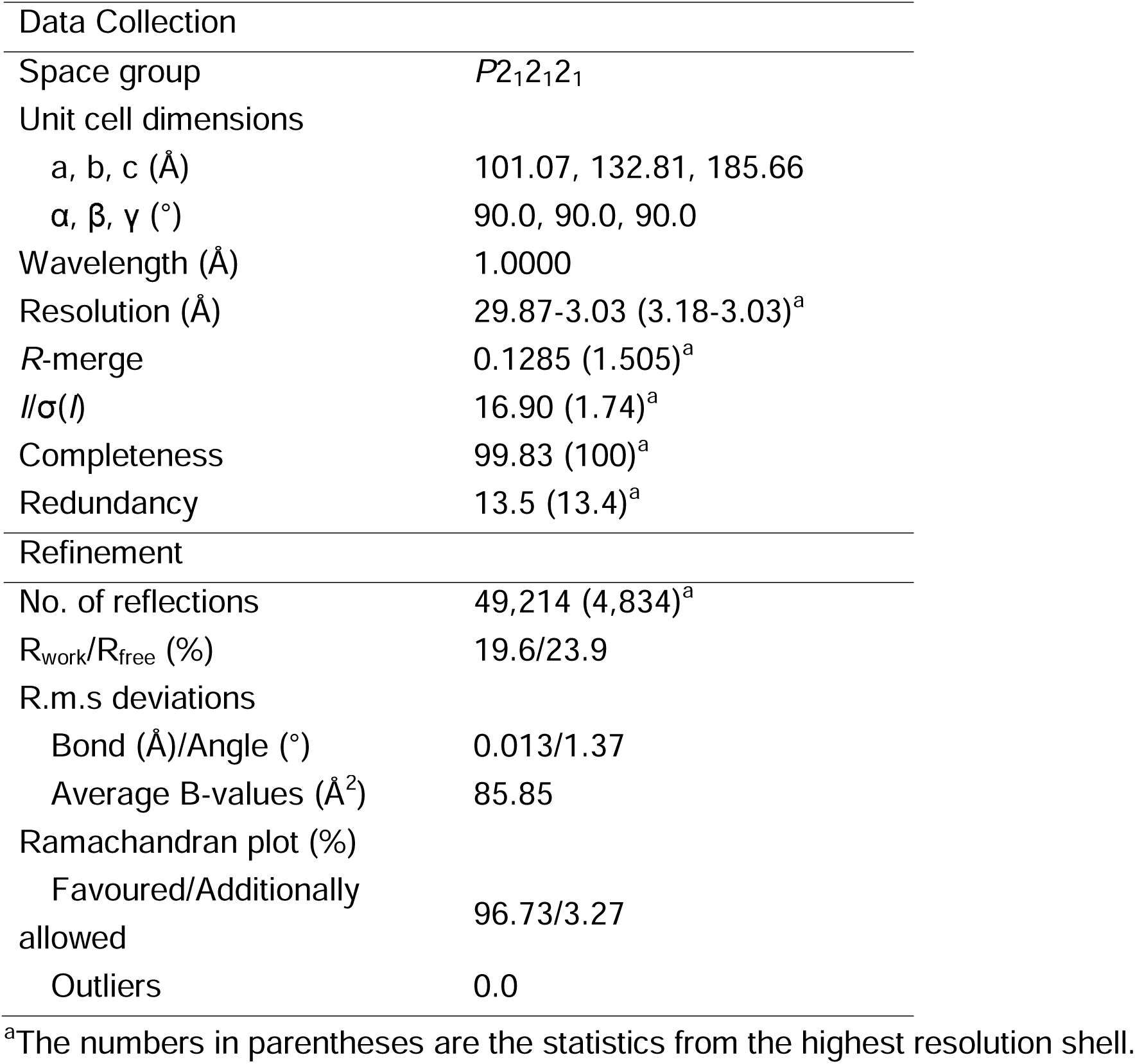
Data collection and refinement statistics.

Typically, the framework regions of V_H_ and V_L_ create a hydrophobic interface between the two domains, and Q2L antibody preserves the conserved hydrophobic residues (Tyr36, Phe98 of V_L_ and Leu45, Phe103 of V_H_) that contribute to this interface formation. We therefore posit that the V_H_ and V_L_ domains of this antibody associate with each other in isolation but dissociate from each other upon binding to RMF/HLA- A*02:01 through V_H_, enabling a unique binding mode that would otherwise be impossible due to severe steric crash between V_L_ and RMF/HLA-A*02:01. The observed unusual structural features demonstrate that the V_H_ domain of this antibody alone provides tight interactions with the pMHC. Two additional structural features stand out in the V_H_ interaction with RMF/HLA-A*02:01. First, V_H_ engages all nine residues of the RMF peptide while making limited contacts with only six HLA-A*02:01 residues (Figure 1B, Bottom). Second, the indole ring of Trp47 is positioned atop one of the “wall-forming” α-helix (α2) of the HLA-A*02:01, suggesting a potential anchoring role for pMHC docking (Figure 1C). Trp47 is a highly conserved framework residue located on a β-strand in a common structural position within the V_H_ domain. We postulated that this docking configuration could be recapitulated by a sdAb. If so, this sdAb and its docking position could serve as a common starting point for discovering TCR-like sdAbs that target diverse pMHCs primarily through peptide contacts, thereby reducing cross-reactivity with non-target pMHCs.

### Design of a sdAb binding to RMF/HLA-A*02:01

We then explored whether the V_H_ domain of Q2L could be produced in a soluble form and exhibit a similar binding affinity. However, the V_H_ domain (residues 1-115) was expressed in Chinese hamster ovary (CHO) cells as a soluble aggregate. This was in sharp contrast with Q2L’s Fab that could be purified as a homogeneous monomeric form, indicating that the Q2L’s V_H_ alone is unstable. Previously, five point mutations introduced into Trastuzumab yielded a soluble monomeric V_H_ domain (Tra_V_H_), and its crystal structure was reported [28]. We sought to graft the key paratope residues of Q2L V_H_ onto Tra_V_H_. A structural alignment of Tra_V_H_ with Q2L_V_H_/RMF/HLA-A*02:01 showed no backbone clashes with RMF/HLA-A02:01, apart from its CDR3H loop (Figure 2A), suggesting the possibility of converting it into a soluble sdAb binding RMF/HLA-A*02:01. After transplanting the CDR3_H loop, grafting two paratope residues on CDR2H (Y57E, R59L), and introducing a Y33A mutation to relieve steric clashes, the model underwent Rosetta FastRelax [29], with only residue 33 designated for redesign. No substantial backbone shifts were observed, and threonine was most frequently enriched at this position. The final selected design included an additional rational F103E substitution to replace an exposed phenylalanine. This design, referred to as RMFsdAb, contains a total of seven point mutations and a four-residue deletion relative to the original Tra_V_H_ sequence (Figure 2B). Noting that Chai-1 [30] accurately predicts the Q2L_V_H_/RMF/HLA-A02:01 structure, we anticipated reliable modeling of RMFsdAb/RMF/HLA-A02:01. The predicted RMFsdAb/RMF/HLA-A02:01 model closely matched the Q2L_V_H_/RMF/HLA-A02:01 crystal structure, with key paratope residues occupying similar spatial positions (Figure 2C). It is noted that, despite conformational differences in the CDR2H loop between Q2L_V_H_ and Tra_V_H_, the Y57E and R59L substitutions on this loop in RMFsdAb remained compatible with RMF/HLA-A*02:01 (Figure 2C).

**Figure 2.**
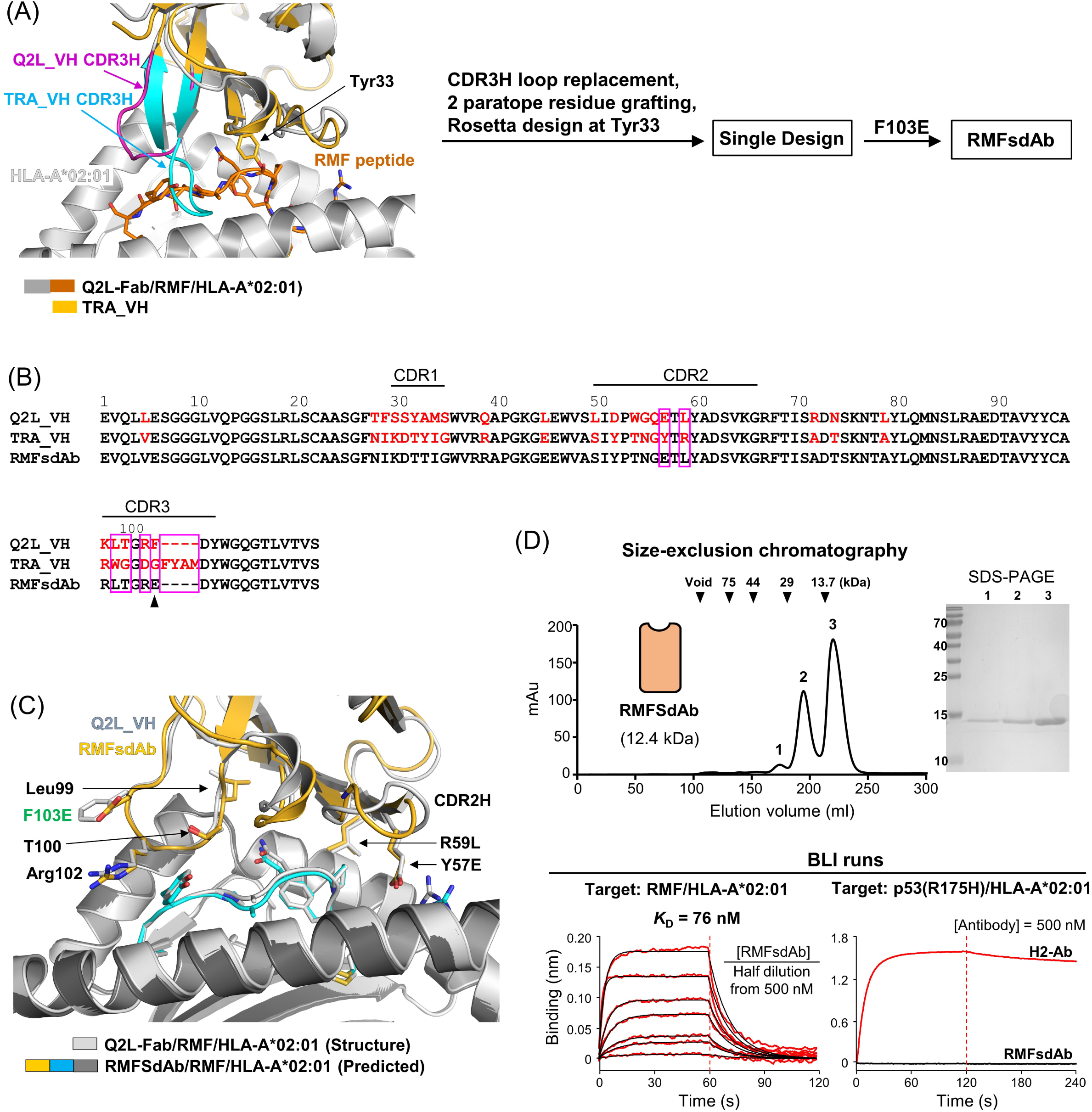
Design, purification and binding affinity measurement of RMFsdAb. (A) Design procedure. Superposition of Tra_V_H_ (PDB entry: 3B9V) onto Q2L-Fab/RMF/HLA-A*02:01 shows overall structural similarity. The CDR3H loops are highlighted in magenta or cyan and labeled. The longer CDR3H loop of Tra_V_H_ clashes with RMF/HLA-A*02:01. Tyr33, which makes a steric clash with the bound peptide, is shown in sticks and labeled. The subsequent design steps are outlined. (B) Point mutations and deletions. The sequences of Q2L_V_H_ and TRA_V_H_ are aligned. Red letters indicate the residues that are different from each other. Pink boxes highlight the five point mutations in RMFsdAb introduced based on the Q2L_V_H_ sequence. The arrowhead marks the rationally replaced F103E mutation to avoid exposed hydrophobicity. (C) Superposition of a predicted structure of RMFsdAb/RMF/HLA-A*02:01 onto the Q2L-Fab/RMF/HLA-A*02:01 structure. The five key paratope residues occupy similar spatial positions (shown in sticks and labeled). Rational F103E mutation is shown and labeled in green letters. The RMF peptide in the predicted structure is shown in cyan. (D) Protein purification and quantification of binding affinity. The elution profile of RMFsdAb from Superdex 75 is shown, along with size marker positions. The inset shows the SDS-PAGE gel of the peak fractions (*Top*). BLI was performed by immobilizing 20 nM RMF/HLA-A*02:01 on streptavidin (SA) sensor tips, incubating with RMFsdAb (monomeric species) and proceeding through the dissociation phase (*Bottom*). In BLI runs with p53^(R175H)^/HLA-A*02:01 immobilized on SA sensor tips, H2-Ab showed binding, whereas RMFsdAb showed no detectable binding.

For experimental validation, RMFsdAb was expressed in *Escherichia coli*. Size-exclusion chromatography showed that approximately 70% of the protein eluted as a monomer, with the remainder as a dimeric species. Biolayer interferometry (BLI) affinity measurement showed the monomeric form bound RMF/HLA-A*02:01 with a dissociation constant (*K*_D_) of 76 nM (Figure 2D). Next, to assess the specificity of RMFsdAb, we prepared HLA-A*02:01 loaded with p53^(R175H)^ peptide (HMTEVVRHC) and an antibody Fab fragment, H2, known to bind this pMHC specifically [7]. Whereas full antibody form of H2 bound strongly to p53^(R175H)^/HLA-A*02:01 In BLI binding assay, RMFsdAb showed no detectable interaction with this pMHC sharing the same HLA subtype (Figure 2D).

### Construction and characterization of HSA D3-RMFsdAb

To address the heterogeneous nature of RMFsdAb (Figure 2D), we fused domain III of human serum albumin (HSA D3; residues 384-585), which is known to be highly soluble and improve the solution behavior of fused client proteins [31, 32]. The resulting fusion protein, HSA D3-RMFsdAb, eluted from a size-exclusion chromatographic column as a single peak, corresponding to a monomeric protein size (Figure 3A). The fusion protein retained a binding affinity for RMF/HLA-A*02:01 comparable to that of RMFsdAb alone (*K*_D_ of 81 nM *versus* 76 nM, respectively; Figure 3A). These results indicate that domain III of HSA greatly improved the solution behavior of RMFsdAb without affecting its binding affinity for RMF/HLA-A02:01.

**Figure 3.**
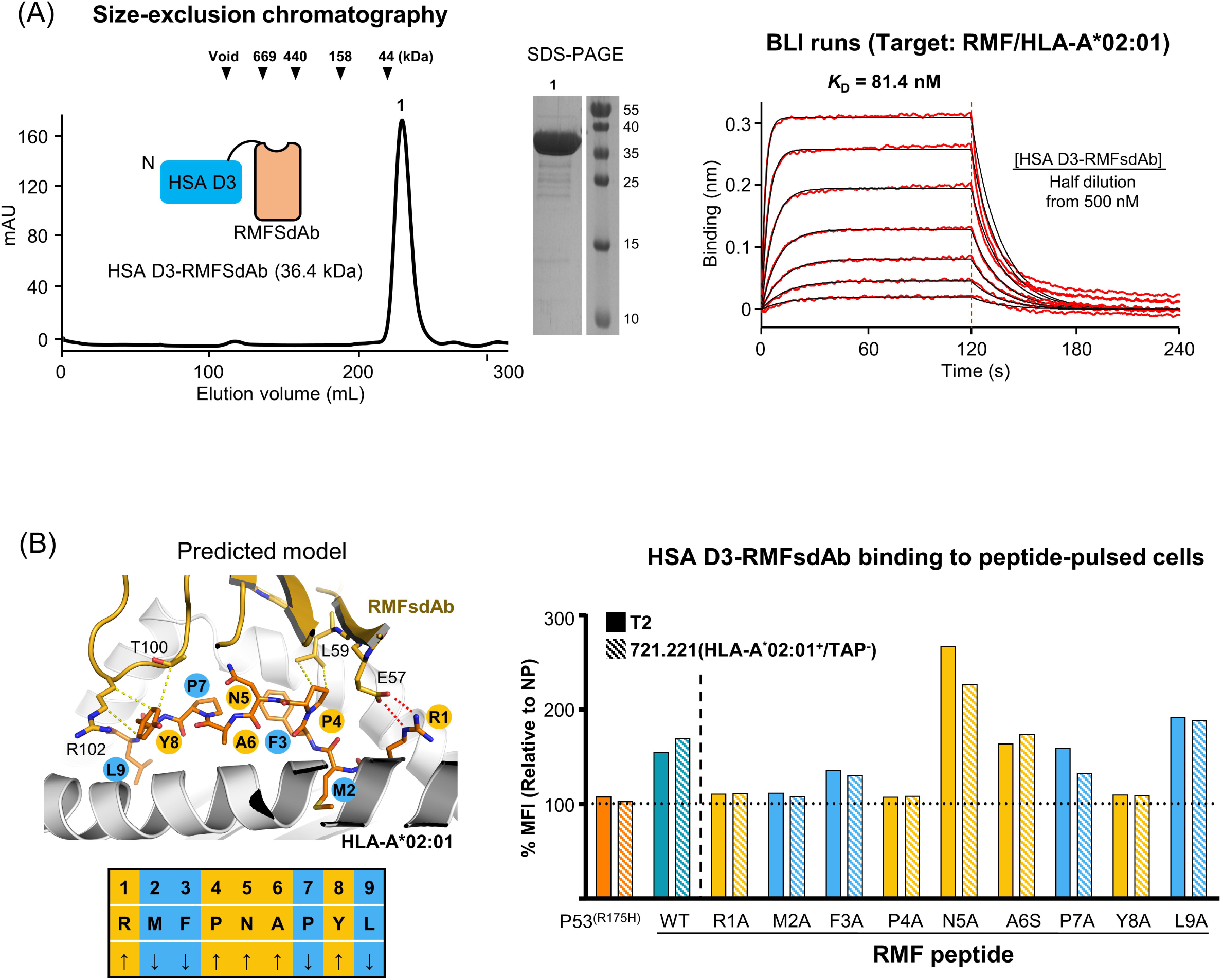
Property of HSA D3-RMFsdAb (A) Purification and binding affinity. A schematic of the HSA D3-RMFsdAb fusion construct is shown. ‘N’ stands for the N-terminus (top). The elution profile from Superdex 200 indicates the monodisperse nature of the fusion protein (*Left*). Binding kinetics were measured by BLI, using SA sensor tips loaded with RMF/HLA-A*02:01, followed by incubation with HSA D3-RMFsdAb and the dissociation phase (*Right*). (B) Alanine scanning of RMF peptide and analysis of binding to cell surface pMHCs. Yellow and red dotted lines indicate key hydrophobic contacts (< 4.1 Å) and ionic bonds, respectively, between the RMF peptide and RMFsdAb in the structural model of RMFsdAb/RMF/HLA-A*02:01. The RMF peptide residues facing RMFSdAb (↑) or HLA (↓) are color-coded in yellow or blue, respectively (*Left*). T2 and 721.221 (HLA-A*02:01^+^/TAP^-^) cells were pulsed with 10 µM of the RMF peptide or the p53^(R175H)^ peptide, and then stained with FITC-labeled HSA D3-RMFsdAb (*Right*). The y-axis represents the mean fluorescence intensity (MFI), expressed as a percentage relative to the unpulsed control (NP).

We next performed cell binding assays with T2 and 721.221 cells to assess the specificity of HSA D3-RMFsdAb for peptide-loaded HLA-A*02:01 on the cell surface. The T2 cell line, derived from a human T cell leukemia line, is naturally deficient in transporter associated with antigen processing (TAP). Because of this deficiency, stable pMHC complexes on cell surface can only form when exogenous peptides are supplied [33]. The 721.221 cell line is a human B lymphoblastoid line that lacks endogenous expression of HLA-A, HLA-B, and HLA-C, but retains expression of TAP [34]; the modified 721.221 (HLA-A*02:01^+^/TAP^-^) cell line stably expresses HLA-A*02:01 but lack TAP expression. T2 cells were pulsed with the RMF peptide, and cells interacting with HSA D3-RMFsdAb were analyzed by flow cytometry. In sharp contrast to unpulsed controls, a clear HSA D3-RMFsdAb–positive population was observed among RMFlllpulsed T2 cells (Figure 3B). In addition, consistent with the BLI binding analysis (Figure 2D), RMFsdAblllHSA did not bind T2 cells pulsed with the p53^(R175H)^ peptide (Figure 3B). These results strongly suggest that HSA D3-RMFsdAb specifically interacts with RMF/HLA-A*02:01 without cross-reactivity to p53^(R175H)^/HLA-A*02:01 and non-specific binding to cell surface components.

To identify RMF peptide residues critical for binding and to assess potential cross-reactivity of HSA D3-RMFsdAb to a similar peptide on the same HLA, cell-binding experiments were performed using RMF peptides containing a positional alanine substitutions (RMF_1A to RMF_9A, except RMF_6S with an Ala-to-Ser substitution).

The p53^(R175H)^ peptide was used as a control. Flow cytometry analysis revealed that substitutions at Arg1, Met2, Pro4, or Tyr8 significantly diminished HSA D3-RMFsdAb binding, indicating that these residues are important for recognition (Figure 3B).

Consistently, in the predicted structure of RMFsdAb/RMF/HLA-A*02:01, these peptide residues, except Met2, engage in hydrophobic or ionic interactions with RMFsdAb (Figure 3B). In the Q2L Fab/RMF/HLA-A*02:01 structure, Met2 is deeply buried within the HLA groove, interacting with Tyr7, Phe9, Met45, Val67, His70, and Tyr99 of HLA-A*02:01. Although Met2 is inaccessible to the antibody, the M2A substitution likely disrupts the peptide binding to HLA-A*02:01, thereby indirectly abolishing HSA D3-RMFsdAb interaction. We note that alanine substitutions at Asn5 and Leu9 enhanced RMFsdAb-HAS-binding compared to the original peptide, likely by altering the overall peptide–HLA-A02:01 interaction in a manner that favors antibody recognition.

### Activation of HSA D3-RMFsdAb CAR-T cells

To assess whether HSA D3-RMFsdAb can activate effector T cells in a CAR format, we engineered Jurkat T cells to express a CAR incorporating HSA D3-RMFsdAb (Figure 4A). These CAR-T cells, designated as HSA D3-RMFsdAb CAR-T cells, were co-cultured with either T2 cells pulsed with the RMF peptide or p53^(R175H)^ peptide, and subsequently IL-2 production was measured by ELISA. The RMF peptide induced a concentration-dependent increase in IL-2 starting at 10 nM (Figure 4B), whereas the p53^(R175H)^ peptide did not elicit IL-2 production even at 10 μM. To further evaluate specificity of activation, we repeated the experiment with 721.221 (HLA-A*02:01^+^/TAP^-^) cells pulsed with the RMF peptide. Similar to the T2 cells, these cells produced IL-2 in a concentration-dependent manner when pulsed with the RMF peptide but not with the p53^(R175H)^ peptide (Figure 4B). Together, these results demonstrate that the HSA D3-RMFsdAb CAR-T cells are selectively activated through specific recognition of the RMF/HLA-A*02:01 complex on the target cells.

**Figure 4.**
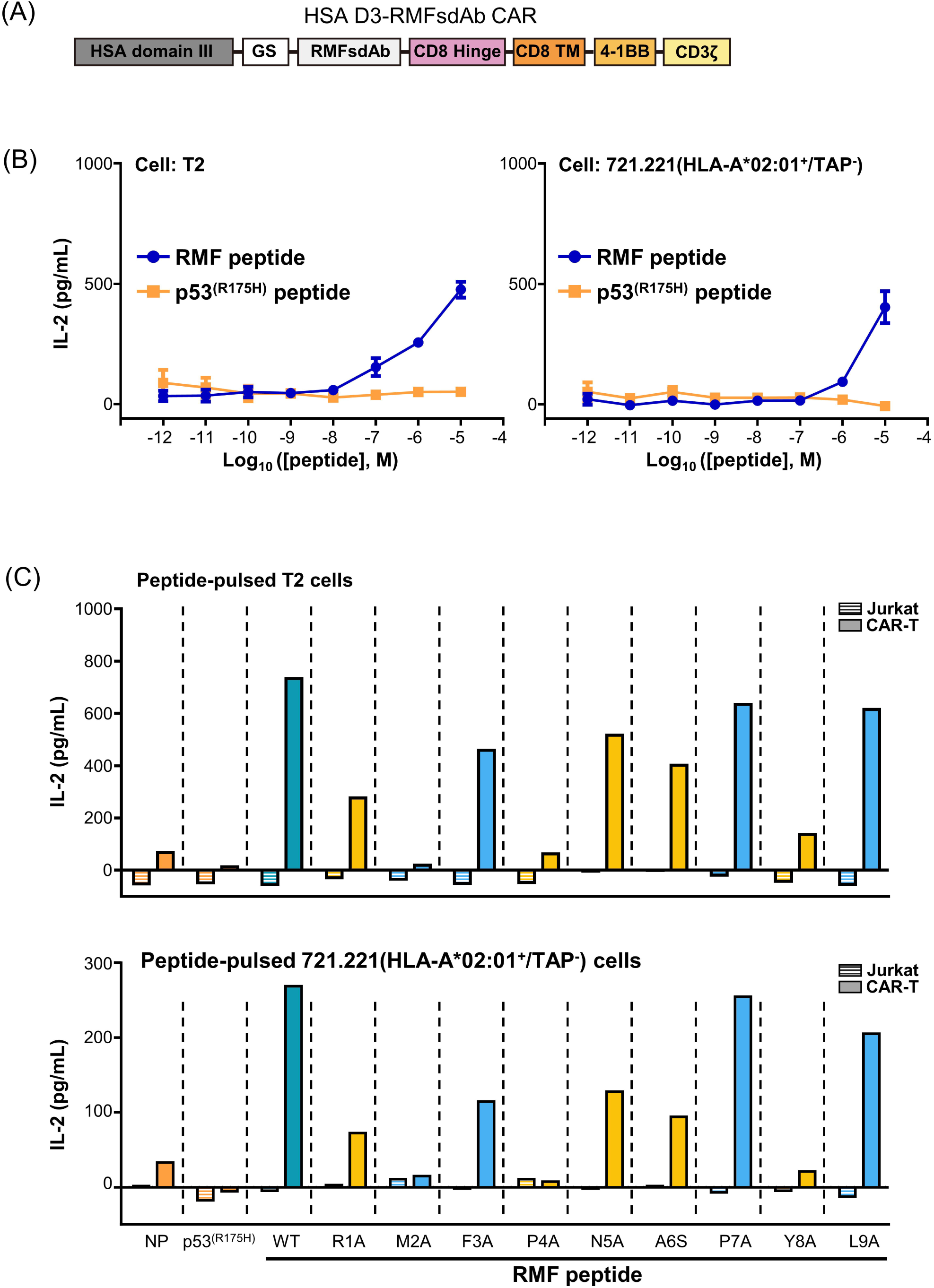
Activation of Jurkat T cells expressing HSA D3-RMFsdAb CAR (A) Schematic of the CAR construct. GS stands for GlySer sequence, and TM stands for transmembrane region. (B) Activation by the RMF peptide. T2 cells (*Left*) and 721.221 (HLA-A*02:01^+^/TAP^-^) cells (*Right*) were pulsed with various concentrations of RMF or p53^(R175H)^ peptides, and co-incubated with HSA D3-RMFsdAb CAR-T cells at a 1:1 ratio. The graphs show the measurement of IL-2 release as a function of peptide concentration. The data represent the mean ± SD of three biological replicates. (C) Diminished activation by mutated RMF peptides. Top panels show T2 cells and bottom panels show 721.221 (HLA-A*02:01^+^/TAP^-^) cells, each pulsed with 10 µM of the indicated RMF peptides. These cells were co-incubated with HSA D3-RMFsdAb CAR-T cells at an effector cell:target cell ratio of 1:1. Controls included unmodified Jurkat T cells and cells pulsed with the p53^(R175H)^ peptide, while NP indicates unpulsed control.

To evaluate whether the activation of HSA D3-RMFsdAb CAR-T cells correlates with the binding affinity of HSA D3-RMFsdAb for the RMF peptide, we repeated the experiment using T2 cells and 721.221 (HLA-A*02:01^+^/TAP^-^) cells pulsed with a panel of peptides, RMF_1A through RMF_9A and RMF_6S. Unmodified Jurkat T cells used as a control showed no activation by any peptide. All mutant peptides showed reduced activation relative to the wild-type RMF peptide in both cell lines (Figure 4C). The extent of this reduced activation paralleled the qualitative binding affinity of HSA D3-RMFsdAb for RMF/HLA-A*02:01 (Figure 3B), because alanine substitutions at key residue positions (M2A, P4A, Y8A) resulted in significantly decreased activation, with the exception of Arg1Ala. At very low or zero peptide loading, the HSA D3-RMFsdAb CAR-T cells showed no activation on either T2 (which express all class I HLAs) or 721.221 cells, demonstrating that HSA D3-RMFsdAb does not cross react with empty HLAs or other cell surface components.

### A bivalent construct of RMFsdAb-HSA D3-RMFsdAb

To further enhance the binding strength of HSA D3-RMFsdAb, we constructed a bivalent variant of the HSA D3 fusion. In this construct, an RMFsdAb moiety was attached to both the N-and C-termini of HSA domain III, yielding a bivalent fusion protein denoted RMFsdAb-HSA D3-RMFsdAb. The bivalent fusion protein was expressed well and purified as a monodisperse monomer by size-exclusion chromatography (Figure 5A). This result indicated that adding a second RMFsdAb does not induce aggregation, enabling the construct to maintain a stable, monomeric state.

**Figure 5.**
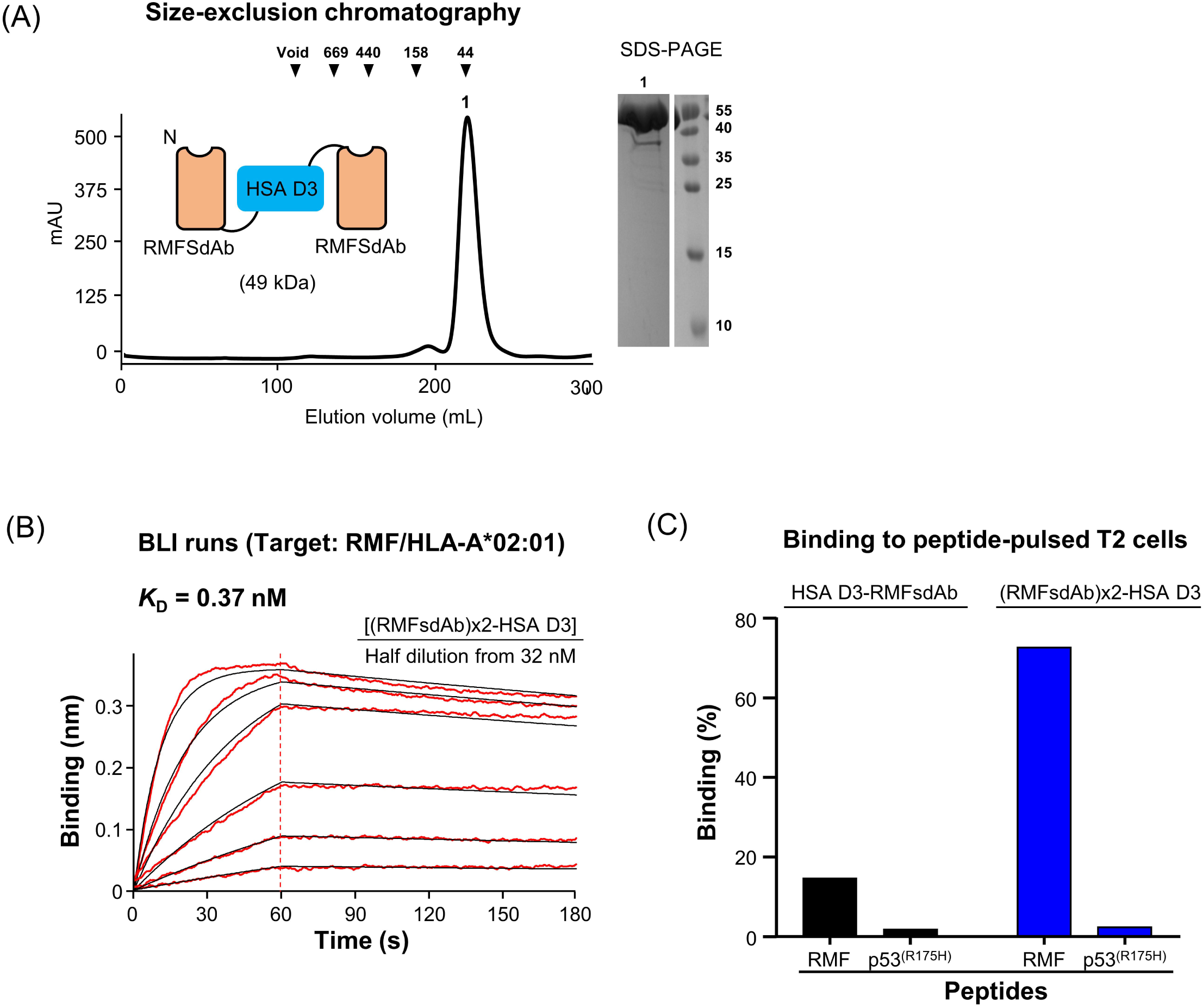
Characterization of HSA D3-RMFsdAb D3-RMFsDAb (A) Schematic and Superdex 75 elution profile of the fusion construct, which was produced and purified as a monomer. (B) Binding avidity measurement. SA sensor tips were loaded with biotinylated RMF/HLA-A*02:01, followed by incubation with the fusion construct and the dissociation phase, yielding a *K*_D_ of 0.37 nM. (C) Compared to the monovalent HSA D3-RMFsdAb, the bivalent construct (indicated as (RMFsdAb)x2-HSA D3) exhibited binding capacity to T2 cells pulsed with the RMF peptide (10 μM), but not to T2 cells pulsed with the p53^(R175H)^ peptide (10 μM). The binding (%) indicates the percentages of the cells shifted from the gate assigned for unpulsed control.

We next evaluated whether bivalency would translate to improved target engagement. BLI measurements revealed that the bivalent RMFsdAb-HSA D3-RMFsdAb exhibited a dramatically enhanced apparent affinity for RMF/HLA-A*02:01, with a *K*_D_ of 0.37 nM (Figure 5B). This represents an ∼200-fold increase in target binding relative to the monovalent HSA D3-RMFsdAb, consistent with the expected avidity gain from dual antigen-binding domains. In agreement with the kinetic data, flow cytometry showed that the bivalent construct bound more strongly to T2 cells pulsed with the RMF peptide than did the monovalent construct (Figure 5C). Importantly, this enhanced binding was antigen-specific: the bivalent RMFsdAb-HSA D3-RMFsdAb displayed no detectable binding to T2 cells pulsed with the p53^(R175H)^ peptide (Figure 5C). Thus, creating a bivalent format greatly improved target cell engagement through avidity while preserving the high specificity of the monovalent HSA D3-RMFsdAb.

## DISCUSSION

For therapeutic applications, TCR-like antibodies must achieve exceptionally high specificity to minimize off-target pMHC engagement and cytotoxicity against healthy cells. This challenge arises from the conserved nature of MHC structures and the small surface area contributed by the displayed peptide, often making it difficult to isolate binders that are peptide-specific yet tolerant of MHC diversity. This study addresses this challenge by designing a nanobody-format TCR-like antibody that predominantly recognizes the bound peptide and avoids significant interaction with HLA.

The V_H_ domain of Q2L Fab demonstrated peptide-centered binding to RMF/HLA-A02:01, making it an attractive scaffold for conversion into a single-domain format. However, the isolated V_H_ was insoluble. Through computational design, CDR3 transplantation and minimal mutations were introduced into a previously optimized monomeric V_H_ scaffold to discover RMFsdAb that binds RMF/HLA-A02:01 with two-digit nanomolar binding affinity. RMFsdAb fusing to HSA D3 significantly improved solution behavior without compromising binding. Importantly, HSA D3-RMFsdAb, exhibited high specificity in both BLI assays and cell-based assays, failing to bind to a closely related p53 mutant peptide presented on the same HLA subtype. Alanine scanning confirmed the functional relevance of several peptide residues, which aligned well with model-predicted paratope interactions, highlighting the peptide-centric nature of recognition.

When expressed in the CAR format, HSA D3-RMFsdAb triggered selective T cell activation in the presence of RMF peptide-loaded HLA-A*02:01, with no response to p53^(R175H)^ peptide-loaded or unpulsed cells. This supports the potential application of RMFsdAb in engineered T cell therapies and further validates its selectivity and specificity.

A nanobody-based BiTE has demonstrated advantages over scFv-based BiTEs, as the latter often suffer from suboptimal dual targeting due to potential mispairing between V_L_ and V_H_ domains and the necessity for a long connecting linker between the two scFv units [35]. Likewise, RMFsdAb based on the soluble V_H_ scaffold derived from human antibody could reduce the risk of off-target interactions compared to conventional Fabs, and its compact modularity allows straightforward fusion to other proteins or multivalent formats as demonstrated with RMFsdAb-HSA D3-RMFsdAb.

A previously reported TCR-like antibody against p53(R175H)/HLA-A*02:01, when converted into a BiTE format, exhibited a *K*_D_ of 86 nM and effectively induced tumor cell killing and xenograft regression in mice. [7]. HSA D3-RMFsdAb exhibits comparable affinity (*K*_D_ of 81 nM), and its bivalent form showed markedly enhanced target binding (*K*_D_ of 0.37 nM). Given that the RMF peptide is derived from the oncogenic WT1 protein—aberrantly expressed in leukemia and multiple solid tumors—RMFsdAb provides a foundational tool for diverse therapeutic formats, including bispecific antibodies, BiTEs and antibody-drug conjugates (ADCs).

Our approach differs from conventional library screening or *in vitro* evolution by leveraging high-resolution structural data and computational design to construct a TCR-like sdAb. Furthermore, the structural features and design strategy described here can serve as a generalizable blueprint for developing sdAbs against other clinically relevant pMHC targets. In conclusion, this work establishes a peptide-focused, computational design platform for generating highly specific, TCR-like sdAbs in a compact and stable format, expanding the repertoire of molecules available for targeting intracellular antigens presented on MHC I.

## METHODS

### Production of pMHC complexes

The RMF peptide (RMFPNAPYL) and the p53^(R175H)^ control peptide (HMTEVVRHC) were synthesized by GenScrpt (GenScrpt, China). Peptide loaded HLA-A*02:01 complexes were produced by protein refolding according to the reported procedures [36]. In brief, denatured HLA-A*02:01 and β2M were refolded in the presence of the synthetic peptide containing 20 mM Tris-HCl (pH 8.0), 400 mM L-arginine, 2 mM EDTA, 0.5 mM oxidized glutathione, 5 mM reduced glutathione and 0.2 mM PMSF. The refolding reaction was incubated at 18 °C for 48 h, followed by concentration to 7.5 mL and buffer exchange using a PD-10 column equilibrated with a buffer solution containing 30 mM NaCl, 20 mM Tris-HCl (pH 8.0). The resulting solution was further purified by size exclusion chromatography using a HiPrep™ 26/60 Sephacryl® S-300 HR column (Cytiva), immobilized metal affinity chromatography (Ni-NTA), and anion exchange chromatography using a Capto™ HiRes Q 5/50 column (Cytiva). The typical yield of final purified pMHC complexes was approximately 2 mg per 2 L refolding reaction.

### Production of Q2L-Fab and sdAb proteins

DNA fragments encoding the V_H_–C_H1_ domain with a C-terminal 6×His tag and the V_L_–C_L_ domain of Q2L were synthesized (IDT) and cloned into a pTT5-derived expression vector (Addgene). The resulting plasmids were transfected into CHO-S cells (Gibco) at 6 × 10 cells/mL using ExpiFectamine CHO (Gibco). Cells were cultured in ExpiCHO Expression Medium (Gibco) for 10 days. Supernatants were harvested by centrifugation at 12,000 × g for 1 h at 4 °C, diluted 1:1 with PBS, and purified by Ni-NTA affinity chromatography (HisPur Ni-NTA resin, Thermo Scientific), followed by size-exclusion chromatography using a HiLoad 26/60 Superdex 75 column (Cytiva) equilibrated with PBS.

The gene encoding RMFsdAb with a C-terminal 8×His tag was synthesized (IDT) and cloned into the pET-22b(+) vector (Novagen). The plasmid was transformed into *E. coli* BL21 (DE3) RIPL cells (Novagen). Protein expression was induced with 200 μM IPTG at 18 °C overnight. Cells were harvested and resuspended in lysis buffer (150 mM NaCl, 20 mM Tris-HCl, pH 8.0), followed by sonication. The cleared lysate was purified using the same purification procedures as for Q2L-Fab.

### Crystallization, structure determination and refinement

Optimized crystallization conditions for Q2L-Fab/RMF/HLA-A*02:01 were established by hanging-drop vapor diffusion at 20 °C, using 2 μL drops prepared by mixing equal volumes of protein and reservoir solutions. Q2L-Fab/RMF/HLA-A*02:01 crystals were obtained using the protein sample (8 mg/mL) and a reservoir solution composed of 8% (w/v) PEG 5000 MME, 5% (v/v) Tacsimate TM (pH 7.0) and 100 mM HEPES (pH 7.0).

For cryoprotection, the crystals were transferred into the reservoir solution supplemented with 18.25% (v/v) glycerol and subsequently flash-frozen in liquid nitrogen. X-ray diffraction data were collected at beamline 5C of Pohang Accelerator Laboratory (Korea), and processed with HKL2000 [37]. Structures were determined by molecular replacement using the predicted structure of Q2L Fab and the crystal structure of RMF/HLA-A*02:01 (PDB entry: 3PHJ) [38] as probe molecules using PHENIX [39]. Electron densities for the two V_L_ domains were weak and vague unlike those for other parts of the complexes, which indicated that the position of V_L_ in the predicted Q2L Fab structure was incorrect. Isolated V_L_ domain was then used for partial molecular replacement with fixed other parts of the complex using PHENIX. The correctly positioned models were refined using COOT [40] and PHENIX [39].

### Biolayer interferometry

BLI experiments were performed using an Octet R8 system (Sartorius). Biotinylated RMF/HLA-A*02:01 at a concentration of 20 nM was loaded onto SA sensor tips (Sartorius). The loaded sensors were then equilibrated in Kinetics Buffer (Sartorius) for 60 seconds. To measure the *K*_D_, RMFSdAb and its HSA D3-fusion constructs at seven different concentrations were subjected to BLI runs with an association step (for 60 or 120 s) followed by a dissociation step (for 60 or 120 s). Binding kinetics were analyzed using the Octet BLI Analysis 12.2.2.4 software package (Sartorius).

For specificity assessment, biotinylated p53^(R175H)^/HLA-A*02:01 and H2-Ab were used as controls, and the experimental procedure followed the same protocol as that used for affinity determination.

### Computational design

After superposing Tra_V_H_ onto Q2L-Fab/RMF/HLA-A*02:01, the CDR3H loop of Tra_V_H_ (residues Arg98-Tyr109) was replaced with that of Q2L (residues Lys98-Tyr105).

Tra_V_H_ residues corresponding to two CDRH2 paratope residues of Q2L were manually replaced (Y57E, R59L), and a bulky residue clashing with RMF/HLA-A*02:01 was mutated to alanine (Y33A). Following model relaxation, position 33 was redesigned with the Rosetta FastDesign protocol. Finally, a variant with an additional F103E substitution was evaluated using Chai-1 for semi-validation of the design.

### Cell Lines

The human B-lymphoblastoid cell line 721.221 (ATCC CRL-1855), T2 (CRL-1992) and Jurkat T cell (ATCC TIB-152) were cultured in RPMI-1640 medium (Gibco, 72400047) supplemented with 10% fetal bovine serum (FBS; Gibco, 12483020). Cells were maintained at 37 °C in a 5 % CO₂ humidified incubator and passaged every 3-4 days to maintain exponential growth.

### Cell-binding assay by Flow cytometry

Pulsed cells were washed one time with ice-cold FACS Buffer (PBS + 1% BSA) and resuspended at 3.75-5.00 x 10^5^ per 100 µL. Cells were stained with pre-incubated HSA D3-RMFsdAb and FITC Anti-6X His tag® antibody (abcam, ab1206) for an hour at 4°C in the dark. After the incubation, the stained cells were washed two times with the cold FACS buffer. After washing, cells were resuspended in 200 µL of the FACS buffer and analyzed on CytoFLEX SRT cell sorter (Beckman Coulter). Data were analyzed using CytExpert SRT (Beckman Coulter). Binding was quantified as mean fluorescence intensity (MFI) within the live cell gate.

### Peptide pulsing

T2 and 721.221 (HLAlllA*02:01⁺/TAP⁻) cells were washed twice with serumlllfree RPMIlll1640, resuspended at 1-2 ×10^6^ cells/mL, and pulsed with peptides (RMF or RMF variants, 1 pM–10 µM) in the presence of 10 µg/mL human β2M. The cells were transferred to lowlllattachment plates (SPL Life Sciences, 32006) and incubated for 18-22 h at 26 °C in a humidified incubator with 5 % CO₂ prior to subsequent functional assays. Pulsing with the p53^(R175H)^ peptide followed similar procedures.

### CAR-T activation assay

In each well of a 96-well v-bottom plate (SPL, 36296), 2.5 x 10^4^ HSA D3-RMFsdAb CAR-T cells were co-cultured with 2.5 x 10^4^ peptide-pulsed T2 or 721.221 (HLA-A*02:01^+^/TAP^-^) in RPMI-1640 supplemented with 10% FBS. The co-cultures were incubated for 24 h at 37 °C in a humidified incubator with 5% CO₂. Following incubation, the culture supernatant was collected to measure IL-2 secretion using the human IL-2 ELISA Kit (BioLegend, 431815) according to the manufacturer’s instructions. The 3, 3’, 5, 5’ tetramethyl benzidine (TMB) substrate was incubated for 20 min, and absorbance was recorded at 450 nm using a Hidex Sense microplate reader (Hidex).

### Generation of stable cell lines

721.221 (HLAlllA*02:01⁺/TAP⁻) was generated through sequential transduction using lentiviral knock-in and lentivirus mediated CRISPR/cas9 knock-out system. In brief, a gene block encoding HLAlllA*02:01⁺ (IDT) was cloned into pLV-EF1a-IRES-Puro lentiviral vector (Addgene #85132). The pLV_EF1a-HLAlllA*02:01⁺-IRES-Puromycin construct co-transfected with lenti-viral packaging plasmids pMD2.2 (Addgene #12259), pRSV-REV (Addgene #12253), and and pMDL g/p PRE (Addgene #12251) into HEK293T cells (ATCC CRL-3216). At 48 h post-transfection, the viral supernatant was exchanged with RPMI-1640 and concentrated to a final volume of approximately 1 mL using Amicon Ultra centrifugal filters (Millipore, UFC9050). The concentrated viral media was filtered through 0.45 µM PES filter (Sartorius, 16533) before use. For transduction, 721.221 cells were plated to 6-well plate one day prior to viral transfection. One mL of prepared viral particles was added dropwise to the cells in the presence of 8 µg/mL sequabrene. At 24 hours post-transduction, cells were selected with 5 µg/mL puromycin. Selection media was refreshed every 2–3 days until the cell population reached approximately 1×10 cells. For single-cell clone generation, cells were serially diluted and plated in 96-well plates, then incubated until colony formation. Expression of HLA-A*02:01 was confirmed by flow cytometry using PE-conjugated anti-human HLA-A,B,C antibody (BioLegend, #311406). Following the selection of HLA-A02:01⁺ cells, TAP1 was knocked out using a lentiviral CRISPR/Cas9 system. A guide RNA targeting TAP1 (5’-CGTTGGCCCGAGCATTGATCCGG-3’), located on chromosome 6, was cloned into the lentiCRISPRv2-blast vector (Addgene #98293). HEK293T cells were transfected with this construct to produce viral particles, which were subsequently used to infect HLA-A02:01⁺ 721.221 cells. Transduced cells were selected with 5 µg/mL blasticidin and subjected to single-cell cloning via serial dilution to establish stable knockout lines.

### Generation of HSA D3-RMFsdAb CAR-T cell line

A second-generation CAR construct was generated by fusing the HSA D3 (residues 384-585) with GS linker to the CD8a hinge and transmembrane domains, in sequence with the intracellular signaling domains of 4-1BB and CD3ζ. A truncated LNGFR (ΔLNGFR, CD271) was included as a selection marker. The CAR cassette was cloned into pLV-EF1a-IRES-Puro lentiviral vector by replacing IRES-Puro element with p2a-ΔLNGFR. The lentiviral particles produced by the CAR transfer vectors transduced Jurkat cells. After 72 h of transduction, ΔLNGFR-positive cells were enriched using anti-CD271 MicroBeads and MACS columns (Miltenyi Biotec) according to the manufacturer’s instructions. The CAR expression was confirmed by flow cytometry using CD271(LNGFR)-FITC Antibody (Miltenyi Biotec 130-112-605).

### Data availability

The coordinates of the Q2L-Fab/RMF/HLA-A*02:01 structures has been deposited in the Protein Data Bank (PDB ID: 9VSO) with the conditions of immediate release upon publication.

## Acknowledgement

This study utilized the Beamline 5C at the Pohang Accelerator Laboratory, Republic of Korea. The research was supported by the National Research Foundation of Korea (Grant Numbers. RS-2024-00440039, RS-2024-00508861, RS-2024-00467046) and by the KAIST Convergence Research Institute Operation Program.

## Author contributions

B.-H.O. and B.-S.J. designed and directed the work. B.-S.J. and T.-S.O. further conceptualized the research. B.-H.O. and B.-S.J. performed the computational design of RMFsdAb. T.-S.O., H.L. and Y.-N.B produced and purified pMHCs and antibodies. T.-S.O. performed the crystallographic work. Y.-S.K supported and S.P.A., J.L. performed cell-based assays. B.-H.O., B.S.J., T.S.O, S.P.A. and Y.-S.K. wrote the original draft. All authors reviewed and accepted the manuscript.

## REFERENCES

1. Thibault, P. and C. Perreault, Immunopeptidomics: Reading the Immune Signal That Defines Self From Nonself. Mol Cell Proteomics, 2022. 21(6): p. 100234.

2. Lowe, K.L., et al., Novel TCR-based biologics: mobilising T cells to warm’cold’ tumours. Cancer Treat Rev, 2019. 77: p. 35–43.

3. D’Angelo, S.P., et al., Afamitresgene autoleucel for advanced synovial sarcoma and myxoid round cell liposarcoma (SPEARHEAD-1): an international, open-label, phase 2 trial. Lancet, 2024. 403(10435): p. 1460–1471.

4. Haus-Cohen, M. and Y. Reiter, Harnessing antibody-mediated recognition of the intracellular proteome with T cell receptor-like specificity. Front Immunol, 2024. 15: p. 1486721.

5. Høydahl, L.S., G. Berntzen, and G. Løset, Engineering T-cell receptor-like antibodies for biologics and cell therapy. Curr Opin Biotechnol, 2024. 90: p. 103224.

6. Liu, H., et al., The Future of TCR-like Antibodies in Diagnosis and Potential Application Targets. Curr Mol Med, 2024.

7. Hsiue, E.H., et al., Targeting a neoantigen derived from a common TP53 mutation. Science, 2021. 371(6533).

8. Augsberger, C., et al., Targeting intracellular WT1 in AML with a novel RMF-peptide-MHC-specific T-cell bispecific antibody. Blood, 2021. 138(25): p. 2655–2669.

9. Campillo-Davo, D., S. Anguille, and E. Lion, Trial Watch: Adoptive TCR-Engineered T-Cell Immunotherapy for Acute Myeloid Leukemia. Cancers (Basel), 2021. 13(18).

10. Huang, D., et al., TCR-mimicking STAR conveys superior sensitivity over CAR in targeting tumors with low-density neoantigens. Cell Rep, 2024. 43(11): p. 114949.

11. Zhang, G., et al., Anti-melanoma activity of T cells redirected with a TCR-like chimeric antigen receptor. Sci Rep, 2014. 4: p. 3571.

12. Duan, Z., et al., CAR-T cells based on a TCR mimic nanobody targeting HPV16 E6 exhibit antitumor activity against cervical cancer. Mol Ther Oncol, 2024. 32(4): p. 200892.

13. Yang, X., et al., Facile repurposing of peptide-MHC-restricted antibodies for cancer immunotherapy. Nat Biotechnol, 2023. 41(7): p. 932–943.

14. Du, H., et al., Targeting peptide antigens using a multiallelic MHC I-binding system.

15. Liu, B., et al., Design of high specificity binders for peptide-MHC-I complexes. bioRxiv, 2024.

16. Sugiyama, H., WT1 (Wilms’ tumor gene 1): biology and cancer immunotherapy. Jpn J Clin Oncol, 2010. 40(5): p. 377–87.

17. Li, X., et al., Wilms’ tumour gene 1 (WT1) enhances non-small cell lung cancer malignancy and is inhibited by microRNA-498-5p. BMC Cancer, 2023. 23(1): p. 824.

18. Salvatorelli, L., et al., Immunohistochemical Expression of Wilms’ Tumor 1 Protein in Human Tissues: From Ontogenesis to Neoplastic Tissues. Applied Sciences, 2020. 10(1): p. 40.

19. Keilholz, U., et al., Wilms’ tumour gene 1 (WT1) in human neoplasia. Leukemia, 2005. 19(8): p. 1318–23.

20. Ohminami, H., M. Yasukawa, and S. Fujita, HLA class I-restricted lysis of leukemia cells by a CD8(+) cytotoxic T-lymphocyte clone specific for WT1 peptide. Blood, 2000. 95(1): p. 286–93.

21. Oka, Y., et al., Human cytotoxic T-lymphocyte responses specific for peptides of the wild-type Wilms’ tumor gene (WT1) product. Immunogenetics, 2000. 51(2): p. 99–107.

22. Holland, C.J., et al., Specificity of bispecific T cell receptors and antibodies targeting peptide-HLA. J Clin Invest, 2020. 130(5): p. 2673–2688.

23. Ataie, N., et al., Structure of a TCR-Mimic Antibody with Target Predicts Pharmacogenetics. J Mol Biol, 2016. 428(1): p. 194–205.

24. Zhao, Q., et al., Affinity maturation of T-cell receptor-like antibodies for Wilms tumor 1 peptide greatly enhances therapeutic potential. Leukemia, 2015. 29(11): p. 2238–47.

25. Dao, T., et al., A dual-receptor T-cell platform with Ab-TCR and costimulatory receptor achieves specificity and potency against AML. Blood, 2024. 143(6): p. 507–521.

26. Chapuis, A.G., et al., T cell receptor gene therapy targeting WT1 prevents acute myeloid leukemia relapse post-transplant. Nat Med, 2019. 25(7): p. 1064–1072.

27. Mun, S.S., et al., Dual targeting ovarian cancer by Muc16 CAR T cells secreting a bispecific T cell engager antibody for an intracellular tumor antigen WT1. Cancer Immunol Immunother, 2023. 72(11): p. 3773–3786.

28. Barthelemy, P.A., et al., Comprehensive analysis of the factors contributing to the stability and solubility of autonomous human VH domains. J Biol Chem, 2008. 283(6): p. 3639–3654.

29. Maguire, J.B., et al., Perturbing the energy landscape for improved packing during computational protein design. Proteins, 2021. 89(4): p. 436–449.

30. Discovery, C., et al., Chai-1: Decoding the molecular interactions of life. bioRxiv, 2024: p. 2024.10.10.615955.

31. Kenanova, V.E., et al., Tuning the serum persistence of human serum albumin domain III:diabody fusion proteins. Protein Eng Des Sel, 2010. 23(10): p. 789–98.

32. Zhao, S., et al., Extending the serum half-life of G-CSF via fusion with the domain III of human serum albumin. Biomed Res Int, 2013. 2013: p. 107238.

33. Henderson, R.A., et al., HLA-A2.1-associated peptides from a mutant cell line: a second pathway of antigen presentation. Science, 1992. 255(5049): p. 1264–6.

34. Shimizu, Y., et al., Transfer and expression of three cloned human non-HLA-A,B,C class I major histocompatibility complex genes in mutant lymphoblastoid cells. Proc Natl Acad Sci U S A, 1988. 85(1): p. 227–31.

35. Yang, X.M., et al., Nanobody-based bispecific T-cell engager (Nb-BiTE): a new platform for enhanced T-cell immunotherapy. Signal Transduct Target Ther, 2023. 8(1): p. 328.

36. Garboczi, D.N., D.T. Hung, and D.C. Wiley, HLA-A2-peptide complexes: refolding and crystallization of molecules expressed in Escherichia coli and complexed with single antigenic peptides. Proc Natl Acad Sci U S A, 1992. 89(8): p. 3429–33.

37. Otwinowski, Z. and W. Minor, Processing of X-ray diffraction data collected in oscillation mode. Methods Enzymol, 1997. 276: p. 307–26.

38. Borbulevych, O.Y., P. Do, and B.M. Baker, Structures of native and affinity-enhanced WT1 epitopes bound to HLA-A*0201: implications for WT1-based cancer therapeutics. Mol Immunol, 2010. 47(15): p. 2519–24.

39. Adams, P.D., et al., PHENIX: building new software for automated crystallographic structure determination. Acta Crystallogr D Biol Crystallogr, 2002. 58(Pt 11): p. 1948–54.

40. Emsley, P., et al., Features and development of Coot. Acta Crystallogr D Biol Crystallogr, 2010. 66(Pt 4): p. 486–501.

